# Microbial signalling of Rhizobium bacteria as a method for cereal crop germination enhancement

**DOI:** 10.1101/788125

**Authors:** Ciara Judge, Emer Hickey

## Abstract

*Rhizobium* nitrogen-fixing bacteria are a well studied microorganism family in the scientific community and it’s effects have been investigated thoroughly since its initial identification. This study aimed to research further the interactions of this bacteria with members of non-legume plant families, i.e. the *poaceae* grass species. This multi-year experimental series focuses on the germination stage of these plants, with varying treatments being applied to the seeds pre-germination in order to first test for a quantifiable effect taking place, and later to characterise the mechanism due to which an effect was noted. The identifier used to establish the presence of an effect was the germination rate of the seeds, which was found by examining the testa of each seed for the emergence of a radicle or plumule. Approximately 15,000 samples were tested during this process, the results of which were subsequently statistically analysed at a 95% confidence level. A conclusive increase of 40% (p<0.0001) was noted in the germination of barley seeds when treated with *R. leguminosarum*. Following tests which confirmed increase enzymatic activity, tests using a different bacterium (*A. tumefaciens*) and further review of existing literature, it was deemed likely that this effect was due to the release of lipo-chitooligosaccharides (LCOs) by the bacteria. It has been established that synthetically isolated LCO’s can positively effect barley germination. It is believed that this is the first time it has been demonstrated that free *Rhizobium* bacteria in suspension can stimulate faster germination in the *poaceae* species due to the natural release of LCO’s by the bacteria.

## 2. Introduction

The following is a summary of an experimental process aimed at determining the effect of various members of the proteobacteria *rhizobia* genus of gram-negative soil bacterium on members of the *poaceae* plant species.

Rhizobium is widely known to be a nitrogen fixing bacterium which forms a symbiotic relationship with members of the legume plant family. (*Ref.1)* The mechanism of rhizobium entry to a plant and nodule formation is characterised by the ability to produce the chemical signalling molecule lipo-chitooligosaccharide (commonly referred to as ‘nod factors’ or ‘LCO’s’).

LCO production is triggered in certain strains of Rhizobia by the presence of flavonoids in the roots of leguminous plants. LCO’s bind to plant receptors, stimulating a response which triggers the nitrogen fixation process. *(Ref.2)* However, LCO’s have also been found to increase the rate of germination. Previous studies have shown that synthetically isolated LCO’s can stimulate faster germination in barley *(Ref.3)*, as can isolated gibberellic acids.

Prior research *(Ref.4)* has shown that the *Bouteloua* grass species contains 19 different flavonoid molecules. *Bouteloua* is a plant genus of the *poaceae* family which is the family in question during these experiments.

While no strain of Rhizobium bacteria has been found to interact with mature plants of the poaceae species, there has been little to no research regarding the impact of it’s application during the germination stage. The possibility of a positive effect being noted in cereal crops (members of the poaceae species) has been long ruled out due to the inability of cereals to form root nodules to host the bacteria. The absence of roots and therefore root nodules during the germination stage has caused the use of nitrogen fixing bacteria such as *Rhizobium* to stimulate faster growth to be overlooked, as the effectiveness of the bacteria outside of a host nodule was considered insignificant. There has however been some research in this area, with studies finding a positive effect on the germination of rice when inoculated with *Rhizobium (Ref.9)*, indicating that the impact of this bacterium during the germination stage warrants further investigation.

*Agrobacterium tumefaciens*, also referred to as *Rhizobium Radiobacter* is a rod-shaped soil bacterium used typically in the process of DNA transformation due to the presence of the Ti plasmid *(Ref.5)*. Their relationship with plants is parasitic rather than symbiotic, causing the contraction of crown gall disease by the host plant *(Ref.6)*. Unlike *R. leguminosarum* and *R. japonicum*, no prior evidence was found to suggest that *A. tumefaciens* releases lipo-chitooligosaccharides at any point during their life cycle. *Agrobacterium tumefaciens* was used later during this research during characterisation of the mechanism as a negative control, i.e. used to confirm that the noted effect can be attributed to LCO release by the bacterium.

According to the above research, a hypothesis was established that the presence of flavonoids in the *poaceae* seeds may trigger LCO release in the bacteria when it is applied in high concentration and a purified state to the seeds. These LCOs will subsequently increase the germination rates of the test seeds.

## 3. Materials and Methods

### 3.1 Germination

#### 3.1.1 Preparation of bacteria

An isolated strain of the test bacterium was cultured in yeast mannitol broth (YMB) for approximately 5 days before centrifugation at 4000G for 8 minutes. The reason for YMB removal was to remove the possibility of a negative effect of the high salt content of the broth on the seeds. While barely and wheat in particular are generally accepted as salt tolerant crops, it was important to eliminate the presence of salt as a variable. The resulting bacterial plug was suspended in sterile water, and the concentration of the mixture was found by serial dilution and turbidity determination via nephelometer. When the bacteria in question was to be killed before application, the culture was lysed via sonification for 5 minutes. Variations of concentration were used for each experiment, with these being quoted in each chart affixed to this report.

#### 3.1.2 Preparation of seeds

The seeds in question were sterilised before inoculation by suspension in a 3% hypochlorite solution and rinsing three times with sterile water in order to remove all hypochlorite residue from the seeds. This sterilisation process eliminates the possibility of viable contamination such as foreign bacteria which would cause variation in the results. The seeds were subsequently placed on an agar medium and each inoculated with a known volume of the relevant treatment (bacteria, control, liquid medium etc.). These samples were subsequently placed in an incubator at a set temperature (varied per experiment). Controls of sterile water inoculation were run in conjunction with each experiment.

#### 3.1.3 Treatments

At various stages of the experimental process different treatments were used. Over the course of the experimental period, germination tests were carried out using *A. tumefaciens* [live], *R. leguminosarum* [live], *R. leguminosarum* [dead], *R. japonicum* [live], *R. radiobacter* [live], *R. radiobacter* [dead], Yeast Mannitol Broth and Sterile Water (de novo).

#### 3.1.4 Measurement of Germination

During experiments the seeds were scanned manually at regular time intervals, or hourly for 72 hours by an Epson Perfection V370 scanner, with the readings for germination taken from the resulting photos. Germination was deemed to have occurred with the seed test was broken by a radicle. The automation of the data collection process decreased the time between measurements which increased the accuracy of the results.

### 3.2 Enzyme Analysis

#### 3.2.1 Preparation of seed samples

Using the method described in section 3.1, seeds were prepared and germinated with either *R. Leguminosarum* or water control inoculation. 10 agar plates were used per treatment, with 2 agar plates per test group being removed each day and kept at −8°C so as to pause the germination process until such time as they could be tested. This continued until completion of the experiment at 5 days.

#### 3.2.2 Alpha-amylase quantification

Each test sample of barley seeds was milled so as to produce a fine powder. Each sample consisted of more than 150 seeds. The samples were incubated at 40°C for 15 minutes and filtered through glass fibre filter paper, before being divided in two in order to provide a control (blank) for each sample. 0.2ml of diluted p-Nitrophenyl B-Maltotrioside (substrate) was added to each sample for exactly 10 minutes before a stopping reagent was added. For the blanks, the stopping reagent was added before the substrate. The stopping reagent halted the reaction and produced a yellow colour, the concentration of which indicated the level of alpha-amylase in the sample by corresponding to the level of unblocked nitrophenyl in the sample. The samples were subsequently incubated for 10 minutes at 40°C. The density of the yellow colour in the samples was measured in a spectrophotometer at 400nm. Each of the samples was also compared to a malt control. These measurements allowed the level of alpha-amylase in each sample to be calculated in ceralpha units.

## 4. Results

### 4.1 Germination

#### 4.1.1 Rhizobium Leguminosarum

As shown in Fig.1, R. leguminosarum was proven to decrease time taken for germination of barley in comparison to a water control (Approximately 10 hours, ANOVA p=0.0001). This result was proven again by a numerous follow up experiments, for example the results seen in Fig.2, where a statistically significant decrease of 22 hours (40.0% reduction) in germination time was noted. In Fig.3 a decrease of germination time can be noted from both volumes of R. leguminosarum applied (28.0% and 8.0% reduction for 25ul and 50ul respectively). Both results are statistically significant, however the effect is less pronounced than with previous experiments. The above results were produced by 3 separate laboratories.

**Fig. 1:**
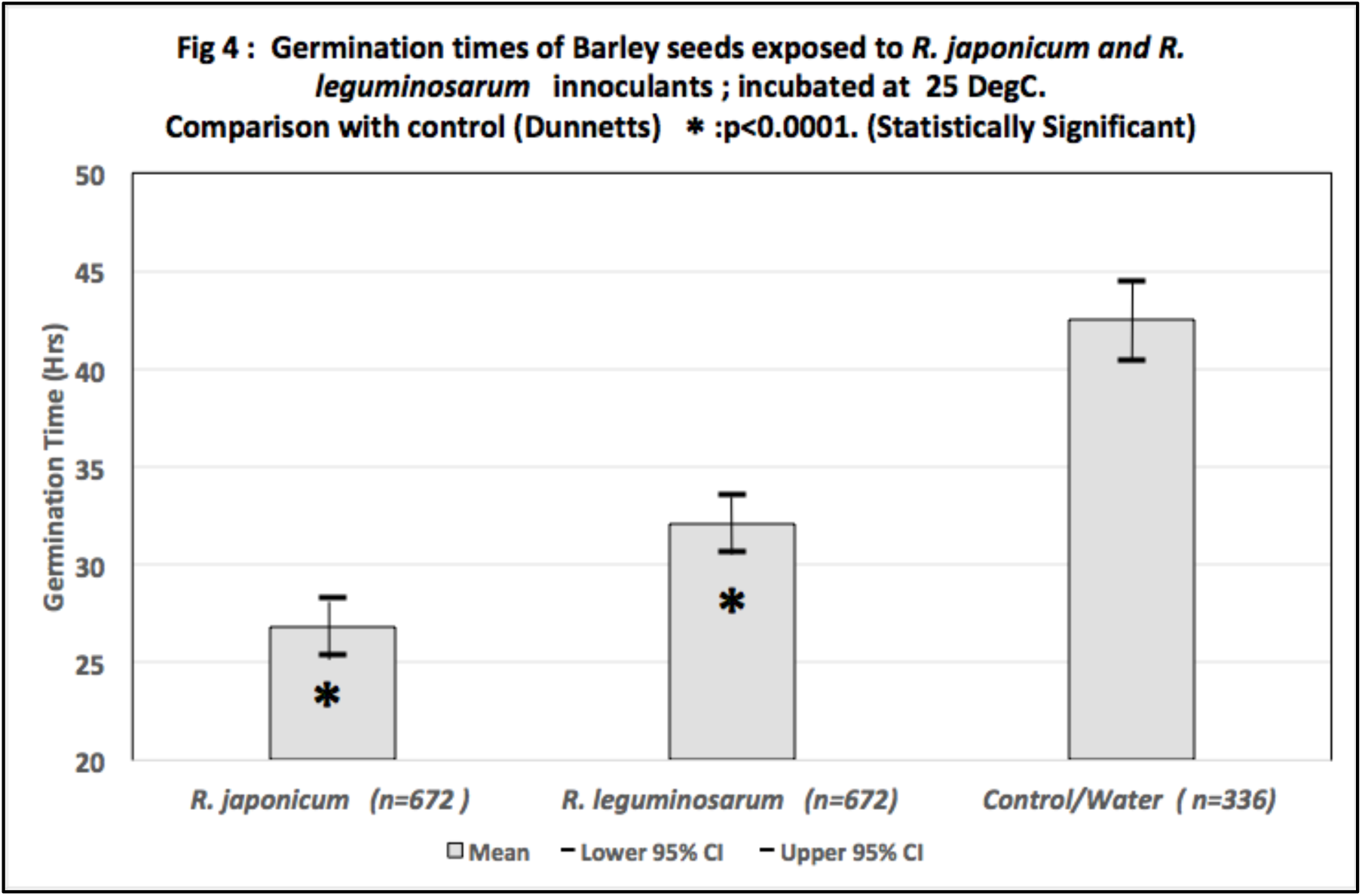
Mean germination time by treatment, Student-t analysis. Inoculum Concentration = [1.6*10^−5^ - 2*10^−7^] cfu/ml

**Fig. 2:**
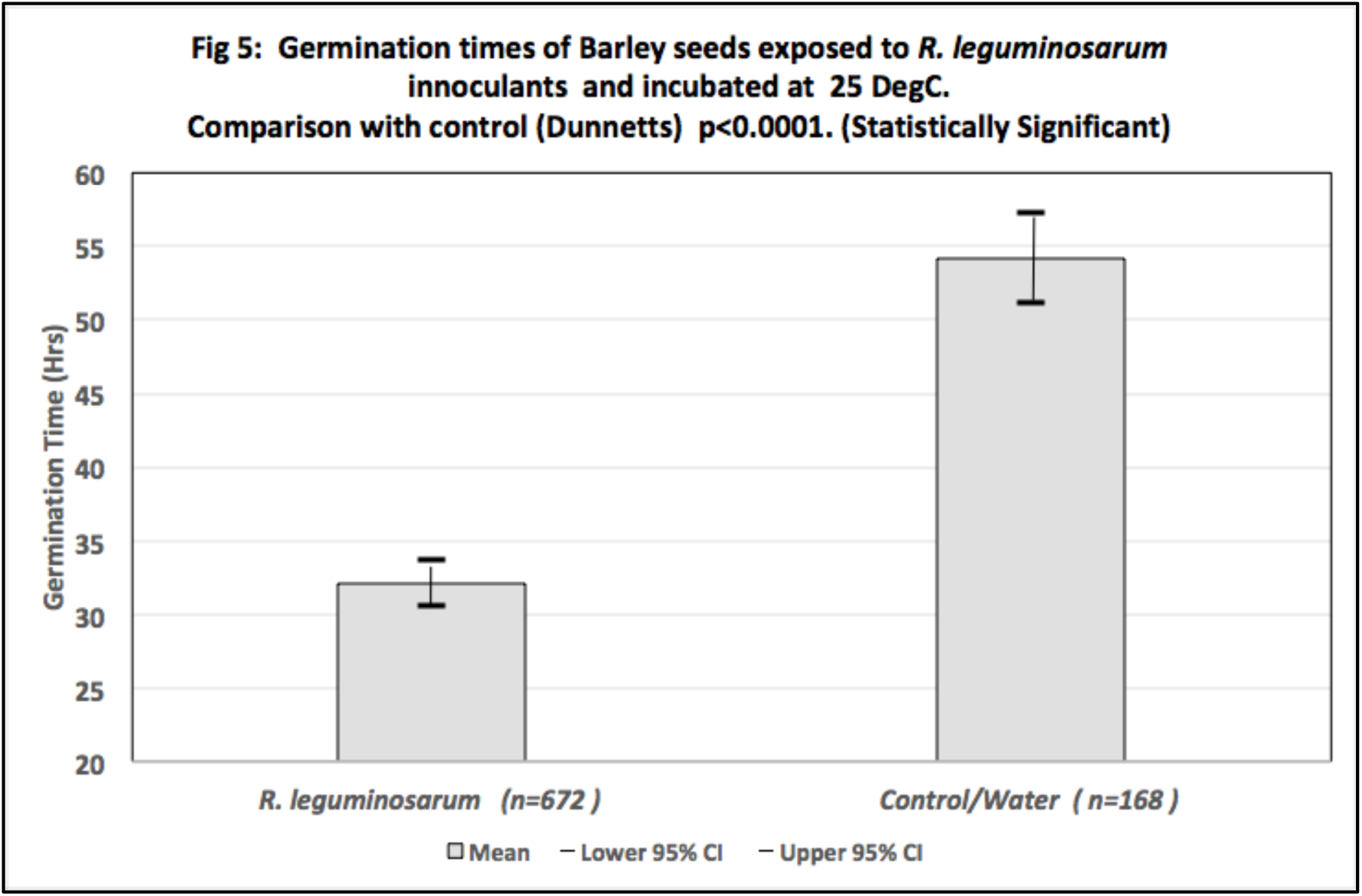
Germination time (hrs) by treatment for R. leguminosarum, Student-t analysis. Inoculum Concentration = [1.6*10^−5^ - 2*10^−7^] cfu/ml

**Fig. 3:**
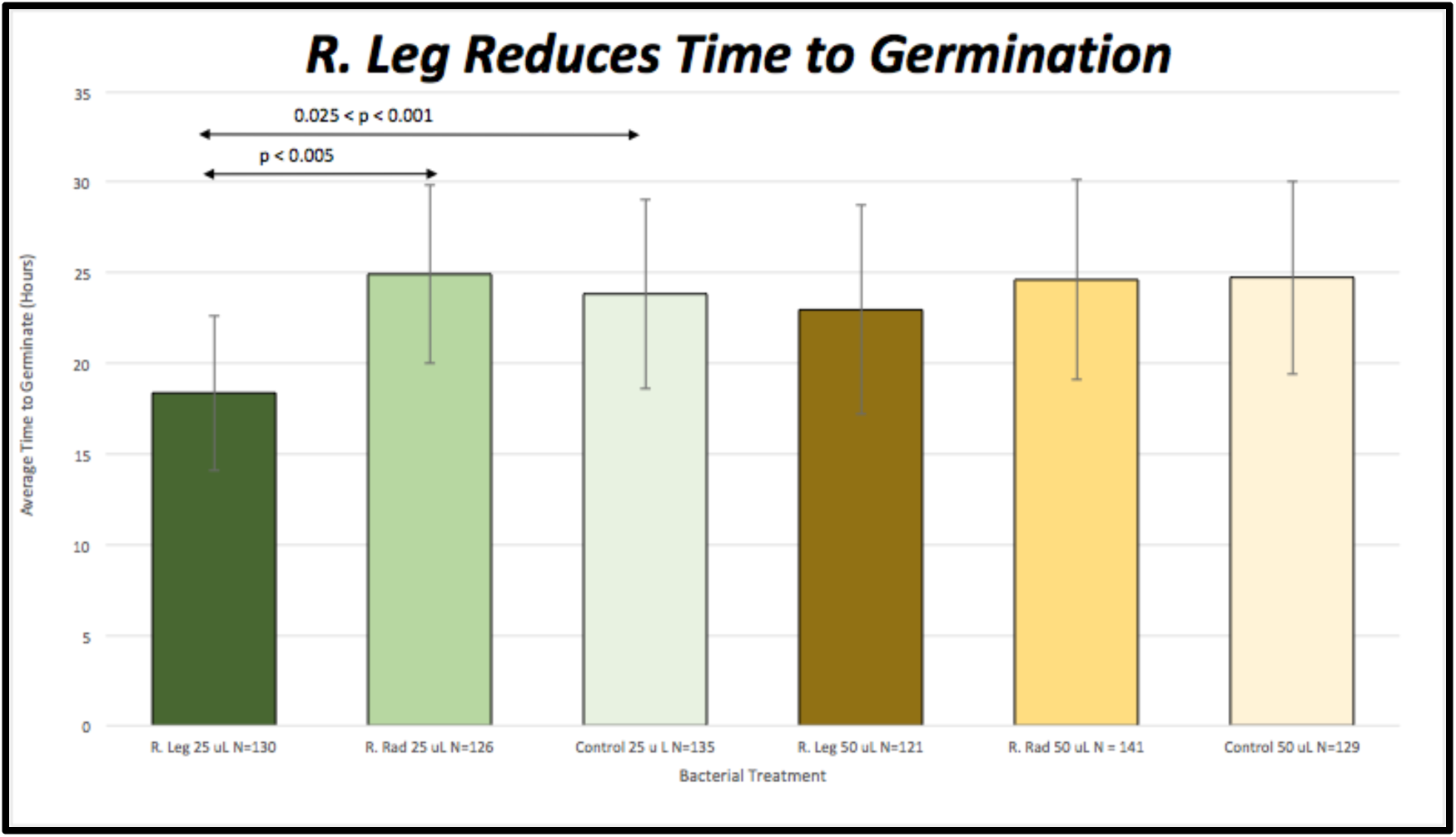
Germination time (hrs) by treatment for R. leguminosarum and A. tumefaciens. Inoculum Concentration = [1.6*10^−5^ - 2*10^−7^] cfu/ml

#### 4.1.2 Rhizobium Japonicum

Referred to earlier in the context of *R. leguminosarum*, Fig.4 shows decreased germination time of barley seeds when treated with *R. japonicum* (12.9% reduction vs water control). R. japonicum also was shown to increase the rate of oat germination in various concentrations (22.3% reduction vs water control)(Fig.4). A concentration curve for this effect could be extracted from the results, however as this was only based on one experimental run, more experimentation is required in this area.

**Fig. 4:**
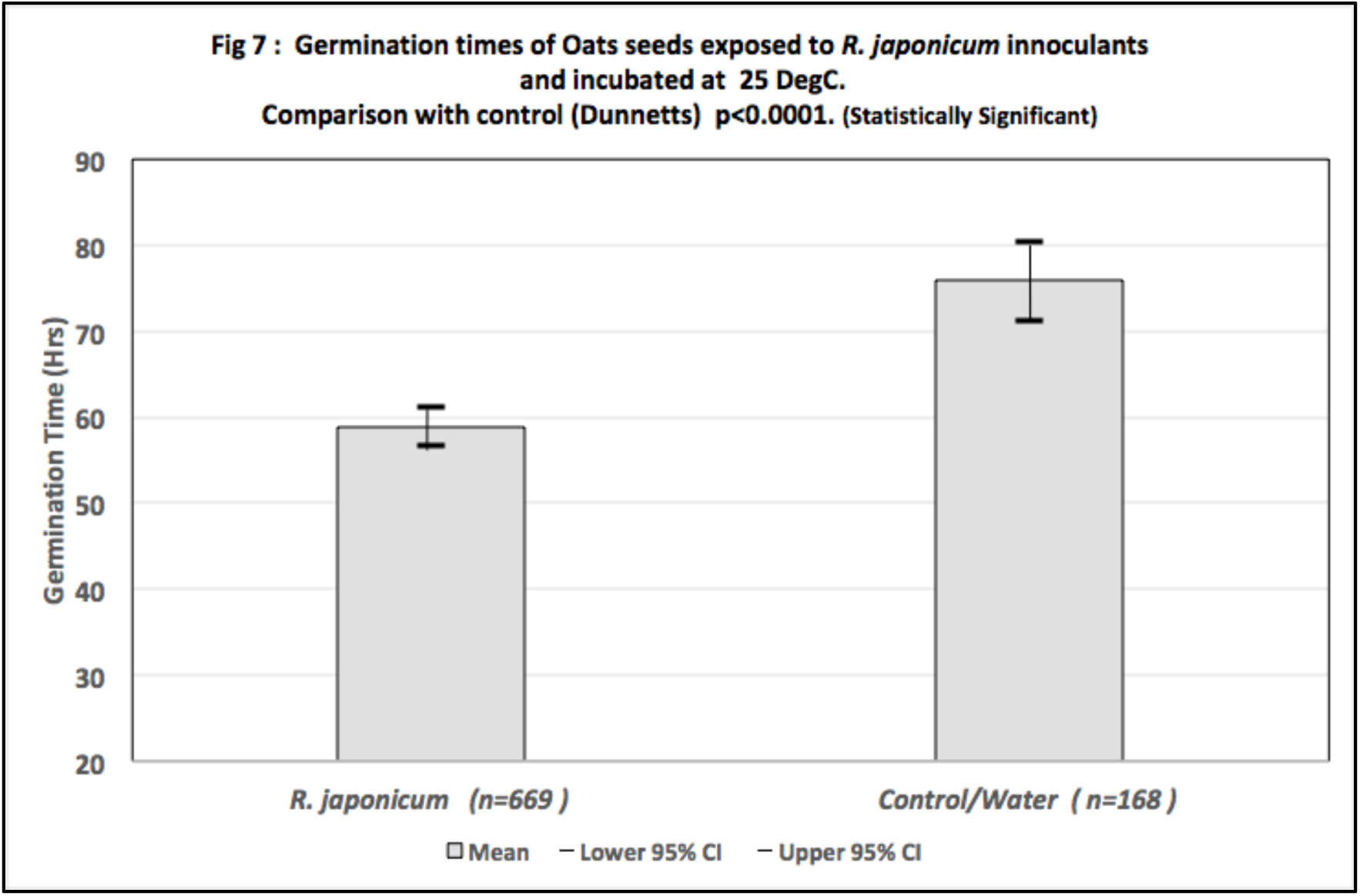
Germination time (hrs) of oat seeds by concentration for R. japonicum. Inoculum Concentration = [1.6*10^−5^ - 2*10^−7^] cfu/ml

#### 4.1.3 Rhizobium Radiobacter (Agrobacterium Tumefaciens)

Agrobacterium tumefaciens was not found to produce any notable or statistically significant effect in any of the experimental runs (Fig.1)

#### 4.1.4 Lysed R. Radiobacter and R. Leguminosarum

No statistically significant effect was noted in seeds treated with non-living *R. radiobacter* and *R. leguminosarum* (Fig.5).

**Fig. 5:**
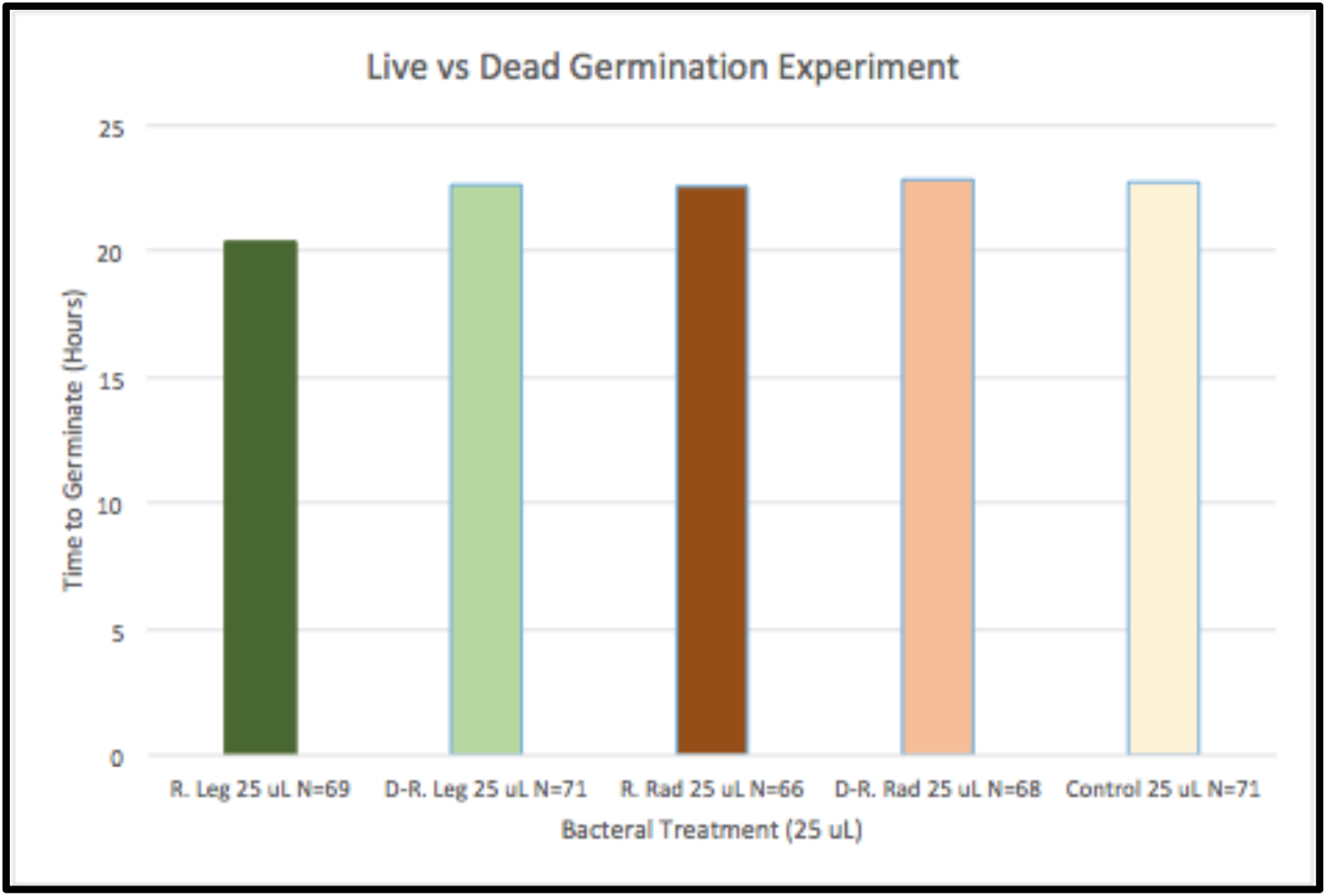
Germination time (hrs) by treatment for live and dead R. leguminosarum and A tumefaciens. Inoculum Concentration = [1.6*10^−5^ - 2*10^−7^] cfu/ml

### 4.2 Enzyme Analysis

Insufficient data points were obtained from the above experimental process to perform a statistical analysis. However, it was possible to track the levels of α-amylase enzyme over time (see Fig.6). Levels of α-amylase are markedly higher in seeds treated by *R. leguminosarum*, although both test samples reached an equal peak level of the enzyme at Day 5 (when most germination had taken place).

**Fig. 6:**
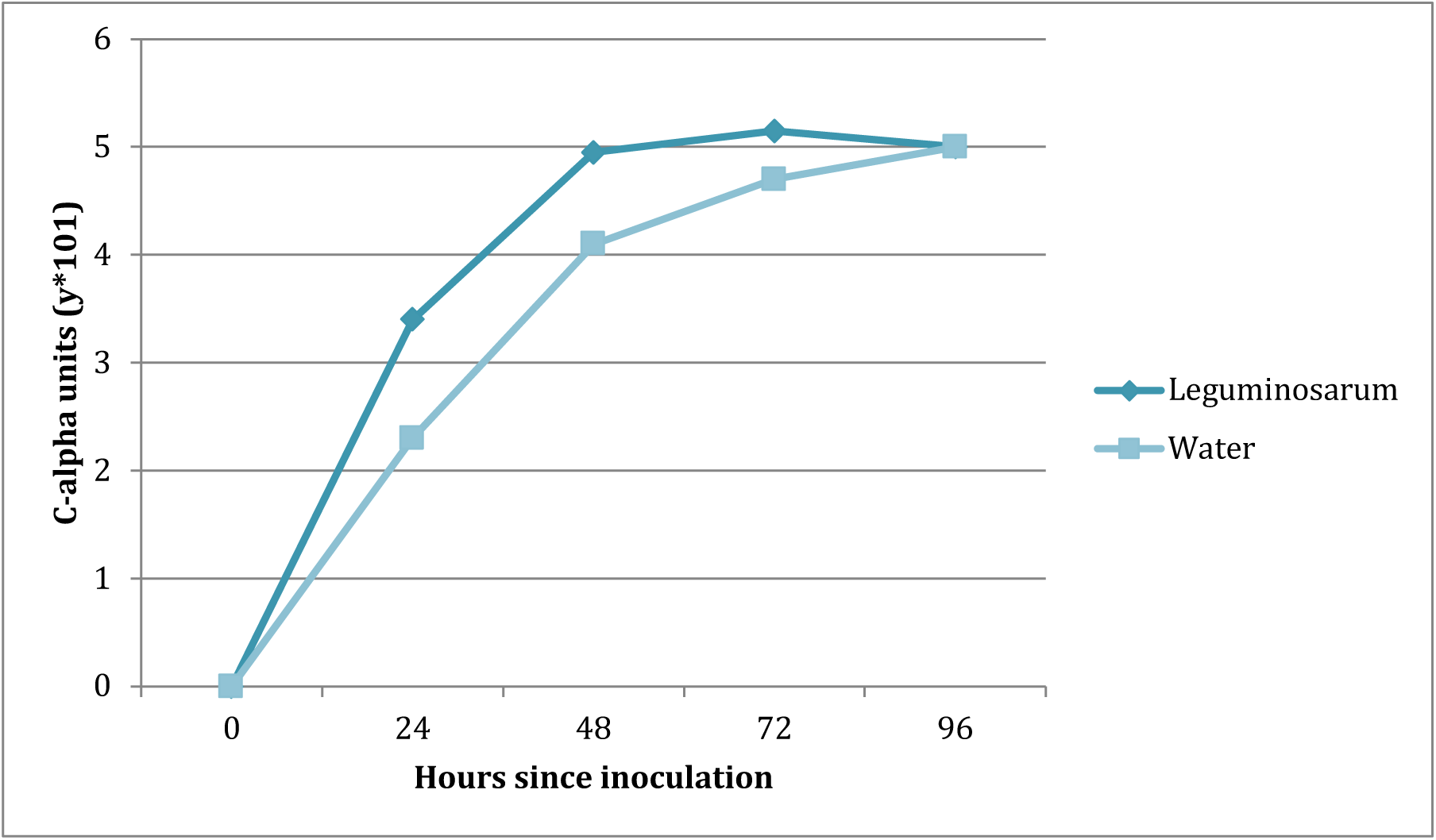
Concentration of alpha-amylase in barley seeds per day for R.leguminosarum. Inoculum Concentration = [1.6*10^−5^ - 2*10^−7^] cfu/ml, Temp. 25°c

## 5. Conclusions

### 5.1 Germination

It can be concluded from the above results that the application of suspensions of the strains *R. leguminosarum* and *R. japonicum* to barley and oat seeds in can produce a notable decrease in the time taken for germination of such seeds. The lack of effect noted in lysed bacteria provides compelling evidence that this effect is triggered by interactions between the live bacteria and seeds, rather than by a physical feature of the bacterial cell.

As seen in Fig.1 and Fig.5, *A. tumefaciens* produced no statistically significant effect. There is no research to support that A. tumefaciens produces lipochito-oligosaccharides, therefore this lack of effect supports the hypothesis that lipochito-oligosaccharides are involved in producing the noted effect. The theory drawn from the above conclusions is that flavonoids in the seeds are triggering the release of LCO’s from the bacteria (*R. leguminosarum* and *R. japonicum*) which in turn mimic the impact of gibberelins and increase the rate of germination.

Due to the fact that live bacteria are required to produce an effect, commercialisation of this process may become more difficult due to handling difficulties. However, as the cultures concerned are sourced from natural soil bacteria this barrier may be easier to overcome *(Ref.11)*.

### 5.2 Enzyme Analysis

The release of the α-amylase hydrolytic enzyme from the aleurone layer is triggered by the presence of active gibberellic acids during germination *(Ref 7)*. As aforementioned, prior research has shown that lipochito-oligosaccharides can act in a similar manner to gibberellins during the germination process *(Ref 3)*. Therefore, while the data set is limited, it is possible that the noted increase in α-amylase levels is cause by LCO’s stimulating earlier and increased production of the enzyme in the seed.

## 6. Discussion

It is believed that the previously described experiments provide evidence of interactions between the chemical signalling molecules of selected members of the *poaceae* grass species (specifically barley) and members of the *rhizobium* family of soil bacteria.

An effect was proven by a very large sample size, with an increased germination rate of up to 40% documented in the test seeds when treated with either *R. japonicum* or *R. leguminosarum*. These results are statistically significant at a 95% confidence level and therefore conclusively indicate a positive impact on germination of the poaceae species.

Furthermore, the lack of effect on the test seeds of bacteria lysed via sonification proves that the previously noted effect was not produced by a physical aspect of the bacteria morphology, but rather a compound or signalling molecule released by live bacteria. It is known from previous research that flavonoids (present in *poaceae* seeds) stimulate the production of LCO signalling molecules from *rhizobium* bacteria *(Ref.8)*, making it likely that these are the compounds in question. This theory is further confirmed by the lack of noted effect on the seeds by *A. tumefaciens*, from which there is no evidence of LCO release at any point in their life cycle *(Ref.10)*.

In research, LCO’s have been proven to act as a substitute for Gibberellic Acids *(Ref.3)*, stimulating the release of hydrolytic enzymes from the aleurone layer of a germinating seed. Therefore an indicator for the presence of LCO’s is heightened traces of hydrolytic enzymes in seeds treated with *rhizobium bacteria*. The marker chosen for this was α-amylase hydrolytic enzyme, and though statistical analysis was not possible, this expected increase was noted in the results of the tests.

A final conclusive test for the above theory would be a quantitative test for LCO release by *rhizobium* bacteria in the presence of the seeds, however the resources for this were outside the scope of this investigation. Therefore, this experiment is a recommended next step from this research. A further recommendation is the repetition of the same experiments, under identical conditions, using seeds which do not contain flavonoids. A lack of noted effect in these experiments would confirm that the effect noted in poaceae seeds was due to LCO release triggered by flavonoids.

Given the above, it is believed that sufficient evidence has been found to support the hypothesis: ***Rhizobium* bacteria can increase germination rates of the *poaceae* grass species due to the release of lipochito-oligosaccharides being triggered by flavonoids present in the *poaceae* seeds. These lipo-oligosaccharides stimulate faster germination by mimicking the effect of gibberellins and triggering increased release of hydrolytic enzymes such as α- amylase from the aleurone layer into the endosperm which ceases seed dormancy**.

## 7. Appendix

### 7.2 Relevant Diagrams

